# An Integrative Study to Investigate Sex-Specific Biomarkers in Bladder Cancer Patients

**DOI:** 10.1101/2024.08.26.609709

**Authors:** Yizhou Wang, Priyanka Bhandary, Kevin Griffin, Jason H. Moore, Xue Li, Zhiping Paul Wang

## Abstract

Bladder cancer shows distinct sex-related patterns, with male patients experiencing significantly higher incidence and female patients facing worse survival outcomes. In this paper, we aimed to address the lack of understanding of the biological mechanisms responsible for this sex-based divergence through an integrative analysis using bladder cancer data from TCGA and GTEx. Our study analyzed various bladder cancer data types, including genomic mutation data, gene expression data, and clinical data. We conducted an in-depth study of protein-protein interactions, pathway analysis, survival analysis, and immune cell correlations. Notably, we discovered that the androgen receptor (AR) related pathways were unique to male hub genes, while the Wnt signaling pathway was unique to female hub genes. Additionally, we identified 14 hub genes with significant sex-biased survival rates, including known DLGAP5, SOX2, LAMA2, and COL5A2, as well as new discoveries of male-specific markers ERCC5, NID1, and ANK2, and female-specific RAD51C, COL22A1 and COL5A2. Furthermore, we identified four male hub genes—DAXX, IKBKB, PDGFRA, and PPARG—that overlapped with immune-related genes. The expression of these genes exhibited differential interactions with immune cells between males and females. These insights could pave the way for more personalized and effective therapeutic interventions tailored to male and female patients.

## Introduction

In recent years, the research community has increasingly focused on the role of sex in cancer, as numerous malignancies exhibit notable differences in incidence and survival rates between males and females [1], [2]. While it is well-established that male and female patients differ genetically due to sex chromosomes and sex hormones, recent studies have revealed that the underlying causal factors extend beyond these apparent distinctions [3], [4], [5]. Consequently, there is a pressing need for more comprehensive investigations into the risk factors and mechanisms driving sex-specific disparities.

Among all cancers, bladder cancer stands out due to its distinct sex-related patterns. Specifically, male patients experience significantly higher incidence rates, whereas female patients face worse outcomes [6]. Although researchers have explored various dimensions, including lifestyle differences, risk factors, and psychological aspects, a critical gap remains related to lack of full understanding of the biological mechanisms responsible for this sex-based divergence [7].

To address this gap, an interrogation from genomic and regulatory perspectives is essential. Our study aims to explore relevant biomarkers, contributing a missing piece to the puzzle of the root causes behind sex-specific variations in bladder cancer. Such information will not only enhance our understanding but also pave the way for tailored precision medicine approaches and informed prognostic strategies in the future.

The exponential growth of publicly available data has revolutionized cancer research, enabling computational investigations. Previous studies have explored The Cancer Genome Atlas (TCGA) data to understand sex differences, ranging from pan-cancer comparisons [8] to specialized bioinformatics tools [9], [10], [11]. However, existing tools often lack the power to systematically compare male and female patients comprehensively. Moreover, sex difference studies in bladder cancer have primarily focused on limited biological aspects, such as gene expression or mutation profiles [12].

Our novel study aims to bridge this gap by integrating multiple layers of information. Specifically, we investigate sex-specific differences in bladder cancer by considering both tumor vs. normal comparisons and male vs. female comparisons. Our comprehensive approach on TCGA data analysis encompasses gene expression patterns, mutation impacts, regulatory factors, and immune effects. By identifying tumor- and sex-specific biomarkers, our study contributes to a better understanding of the biological functions and pathways that relate to sex difference in bladder cancer.

One limitation of TCGA data is its scarcity of normal sample data, which poses a significant challenge for bladder cancer research. This scarcity restricts the use of TCGA data as a reliable control for most tumor types. To address this, researchers have explored alternative control samples from external resources such as the Genotype-Tissue Expression (GTEx) project [13], [14]. GTEx provides diverse non-disease tissue samples, allowing us to augment the TCGA control data. For this reason, the analysis strategy of our study has incorporated GTEx control samples. This approach enables the investigation of cancer-specific biomarkers by comparing tumor samples to these external controls.

In this study, we aim to elucidate the underlying mechanisms driving sex disparities in bladder cancer, by employing an integrative approach, leveraging both GTEx control samples and TCGA data. Our methodology involved a comprehensive comparative analysis of bladder cancer datasets, stratified by sex. Our bioinformatics analyses focused on identifying tumor-specific significant genes, delineating key biomarkers indicative of sexual dimorphism, exploring regulatory factors such as microRNAs and lncRNAs, and elucidating pertinent biological pathways. Furthermore, survival and prognosis analyses were conducted to assess the clinical relevance of identified biomarkers.

## Results

### 1. Mutations in male and female BLCA patients

The mutation profile from the whole-exome sequencing (WXS) data of bladder cancer (BLCA) patients from The Cancer Genome Atlas (TCGA) database was analyzed. Variants were categorized into nine distinct groups based on their impact on protein-coding genes, evaluated separately for male and female samples (Fig. 2A, B). Our analysis demonstrated that the mutation profiles between male and female samples are highly similar, with missense mutations being the predominant variant type. Among these variants, single nucleotide polymorphisms (SNPs) were significantly more prevalent than insertions (INS) and deletions (DEL). Furthermore, the C>T transversion emerged as the primary type of single nucleotide variant (SNV) in BLCA for both male and female patients. Among the top 10 genes with the highest mutation frequency for both male and female patients, the shared genes with the mutation frequency (female, male) were TTN (43%, 46%), TP53 (51%, 46%), KDM6A (28%, 25%), KMT2D (25%, 31%), MUC16 (22%, 30%), PIK3CA (22%, 21%), and ARID1A (19%, 26%). Unique to the top 10 genes in female patients were ELF3 (15%), FAT1 (16%), and HMCN1 (16%), whereas unique to male patients were SYNE1 (22%), KMT2C (19%), and RB1 (19%). Genes exhibiting the top 50 mutation frequencies are illustrated in Supplementary Figure 1A and B.

**Figure 1.**
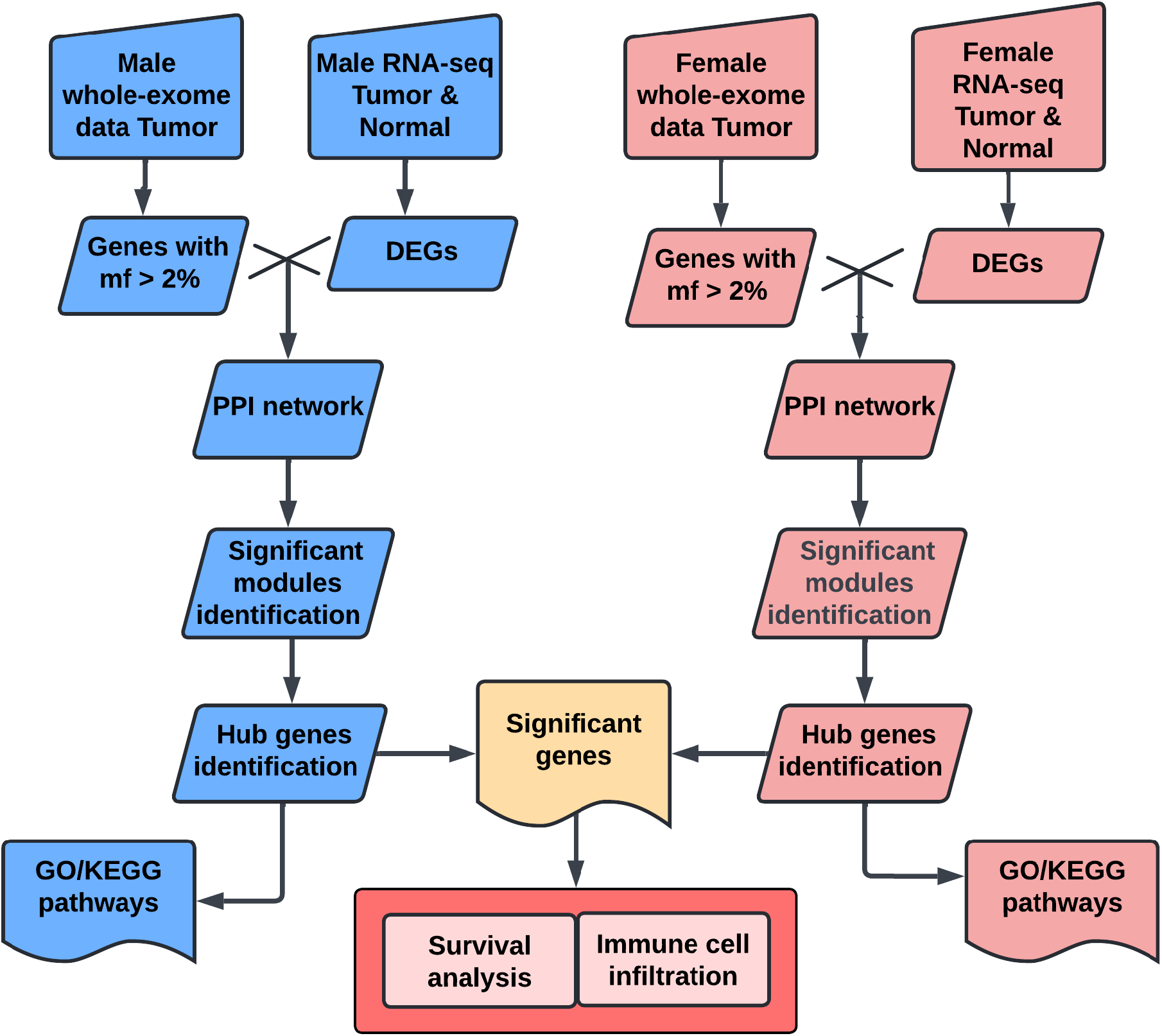
Workflow chart of the bioinformatics analysis performed with TCGA and GTEx BLCA datasets. DEG, differentially expressed gene; GO, Gene Ontology; KEGG, Kyoto Encyclopedia of Genes and Genomes

**Figure 2.**
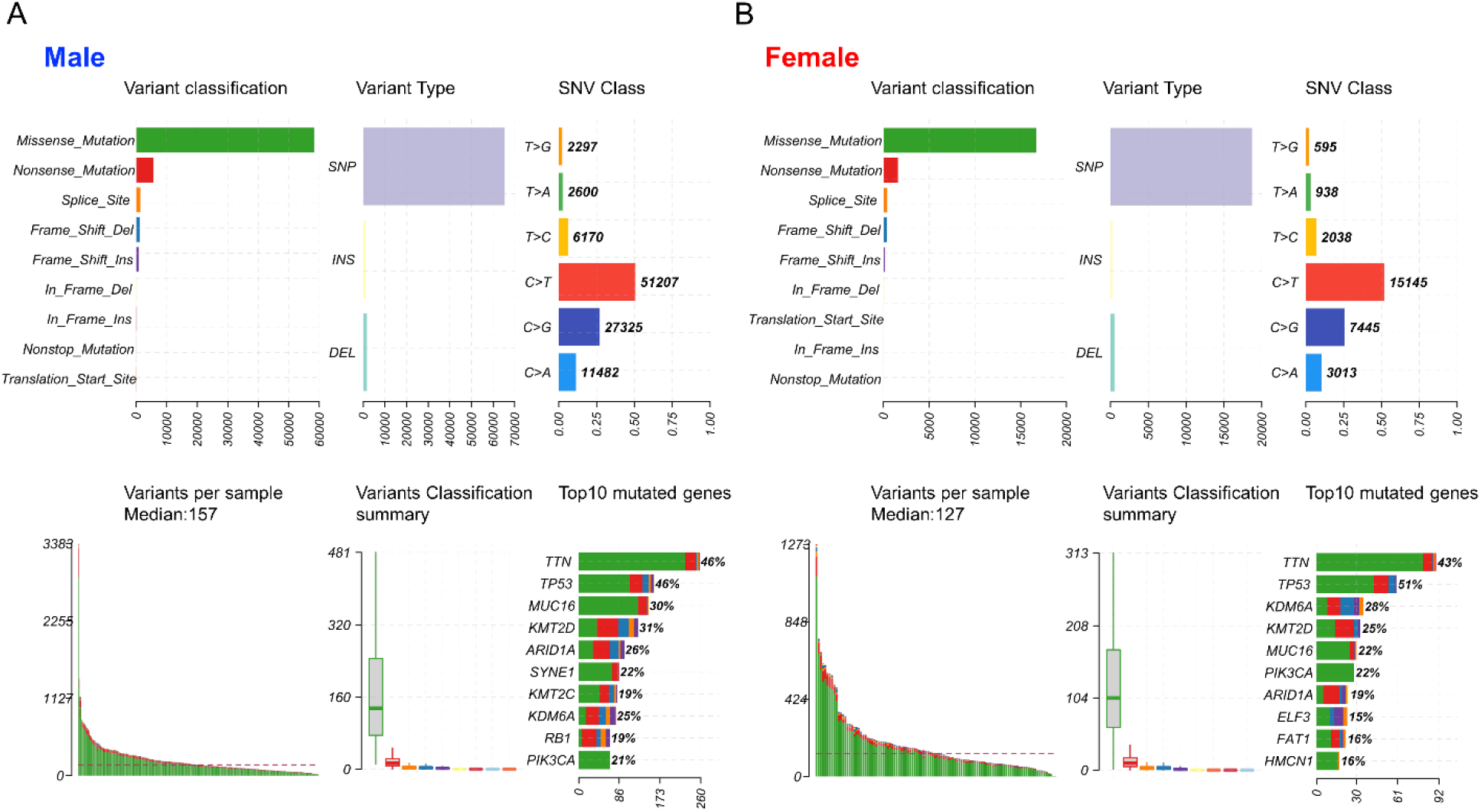
The landscape of mutation profiles in BLCA patients, including variant classifications, variant types, SNV class, variants per sample, variant classification summary and top10 mutated genes. (A) male and (B) female patients.

### 2. Identification of DEGs and overlapped DEGs with intermediate mutation frequency

Gene expression data of BLCA patients for 19,127 protein-coding genes were obtained from the TCGA and GTEx datasets. We identified 7,916 differentially expressed genes (DEGs) in female patients and 11,827 DEGs in male patients when comparing tumors to normal samples. Bidirectional clustering heatmaps indicated significant differences between tumors and normal samples in both sexes (Fig. 3A, B). Genes with mutation frequencies between 2% and 20% in a given type of tumors were classified as having intermediate frequencies [15]. Accordingly, we focused on the DEGs with a mutation frequency greater than 2%. Our analysis identified 871 DEGs in female samples (349 upregulated and 522 downregulated) and 1,666 DEGs in male samples (687 upregulated and 979 downregulated). Venn diagram analysis (Fig. 3C) showed that 415 DEGs were unique to females and 1,210 DEGs were unique to males. Among these, 203 genes were upregulated and 212 were downregulated in female samples, while 541 genes were upregulated and 671 were downregulated in male samples. Additionally, 454 DEGs (146 upregulated and 308 downregulated) were common to both female and male BLCA patients.

**Figure 3.**
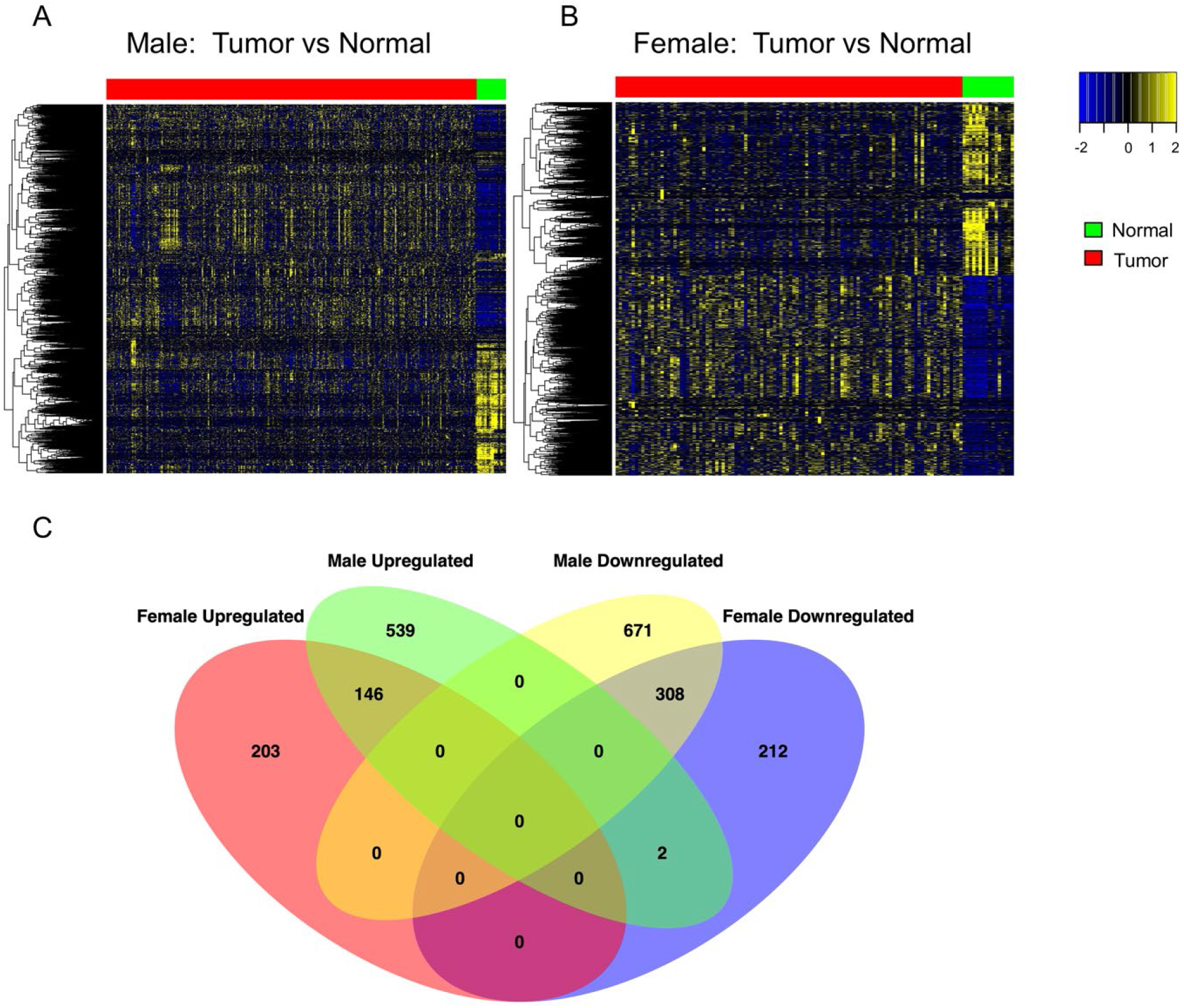
Differential gene expression analysis to identify sex-dependent DEGs. Heatmaps of DEGs in (A) male and (B) female BLCA patients. The top red and green bars indicate tumor bladder tissue and normal bladder tissue, respectively. Blue boxes represent downregulated genes, and red boxes represent upregulated genes. (C) Venn diagram of DEGs with intermediate mutation frequency (>2%) in the two sexes. DEG, differentially expressed genes.

### 3. Protein-protein interactions (PPIs), key module analyses and hub genes in networks

DEGs with intermediate mutation frequencies from both sexes were imported into STRING to analyze PPIs among the various gene targets. PPI networks were constructed using Cytoscape for male and female samples, respectively. DEGs with more than one connection to other DEGs were retained for further analysis. Gene modules were clustered using the MCODE plugin, revealing a total of 32 modules in males, with four modules having scores greater than 10, and 21 modules in females, with two modules having scores greater than 10 (Fig. 4A, B).

**Figure 4.**
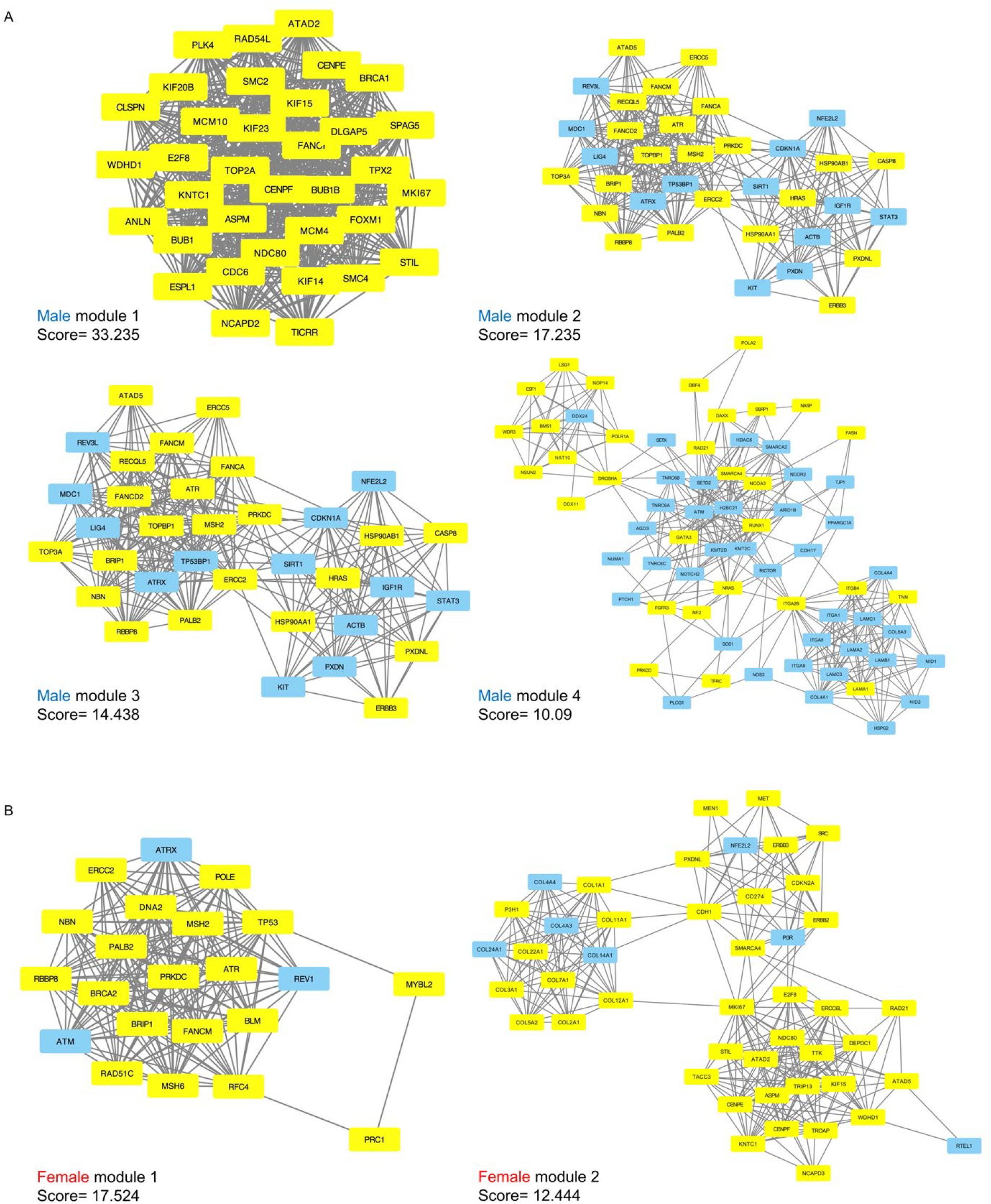
Significant submodules of PPI analysis (Molecular Complex Detection scores > 10). (A) male significant modules (1-4) (B) female significant modules (1-2). Yellow represents upregulated nodes and blue represents downregulated nodes.

In the male samples, module 1 had a score of 33.23 with 35 nodes and 565 edges; module 2 had a score of 17.23 with 35 nodes and 293 edges; module 3 had a score of 14.44 with 65 edges; and module 4 had a score of 10.09 with 68 edges. In the female samples, module 1 had a score of 17.52 with 22 nodes and 184 edges, and module 2 had a score of 12.44 with 46 nodes and 280 edges. From the four male modules, 203 genes were identified, with 117 upregulated and 86 downregulated. From the two female modules, 68 genes were identified, with 58 upregulated and 10 downregulated (Supplementary Table 1). All these genes showed a degree greater than 10 according to cytoHubba, indicating their potential as hub genes.

### 4. Functional and pathway enrichment analysis of sex specific hub genes

We further analyzed the function of the genes in each male and female significant module by GO-BP and KEGG. A total of 1421 significant GO terms for the male hub genes and 558 GO terms for the female hub genes were obtained (adjusted p < 0.05). Several pathways were common to both sexes, while additional pathways were primarily enriched in one sex, including specific pathways enriched in either males or females (Fig. 5A-D, Supplementary Table 2). The top GO terms for both sexes were primarily enriched in “cell cycle”, “DNA replication”, “DNA repair”, “DNA recombination”, “nuclear division”, and “chromosome segregation, separation and organization”, “chromosome organization” and “response to xenobiotic stimulus” (Fig. 5A, B). Male hub genes also demonstrated unique enrichment in the pathways related to the protein kinase activity, basement membrane organization, T cell function related pathways, epigenetic regulation of gene expression and androgen receptor (AR) signaling pathway. Female hub genes showed unique enrichment in the pathways related to the Wnt signaling pathway. For KEGG pathways, both sexes exhibit the primary enrichment in multiple cancer related pathways, such as “p53 signaling pathway”, “PI3K-Akt signaling pathway”, “Fanconi anemia pathway”, “focal adhesion”, “ECM-receptor interaction”. Male hub genes also uniquely enriched in the “FoxO signaling pathway”, “HIF-1 signaling pathway”, “mTOR signaling pathway”, “Renal cell carcinoma” and “Chemokine signaling pathway”, which are all crucial in tumorigenesis (Fig. 5C, D).

**Figure 5.**
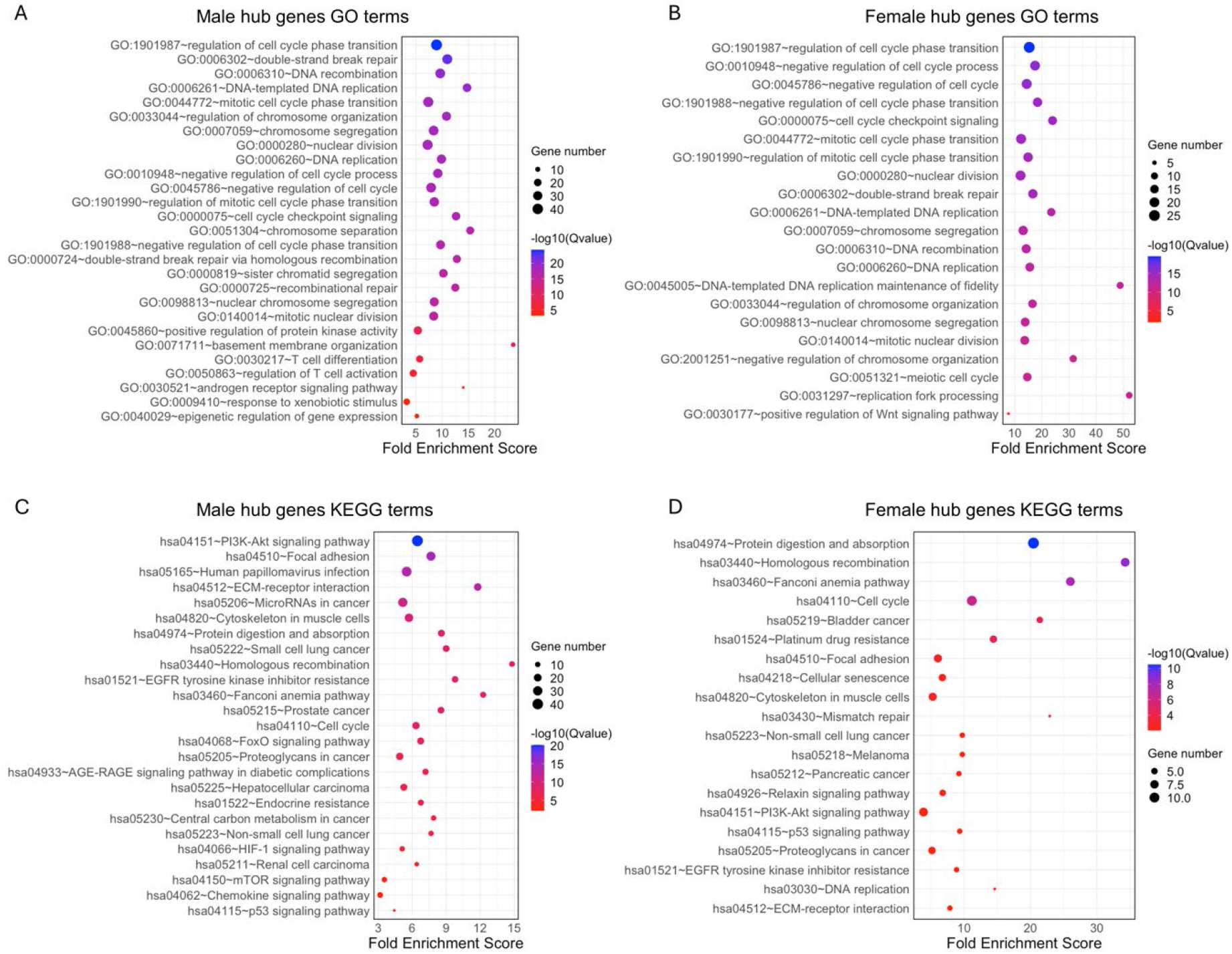
GO-BP terms and KEGG pathway enrichment of genes in key modules of the two sexes. Top 20 GO terms and selected sex specific pathways in (A) male and (B) female patients, respectively. Top 20 KEGG pathways in (C) male and (D) female patients, respectively. GO, Gene Ontology; KEGG, Kyoto Encyclopedia of Genes and Genomes.

### 5. Survival analysis for genes in key modules

We compared the potential hub genes identified in male and female key modules and identified 161 male unique genes, 26 female unique genes and 42 common genes (Fig. 6A). To investigate the effects of each module on patient survival, we analyzed the prognostic value of the hub genes from four significant male and female modules using the Kaplan-Meier algorithm. In the male modules, 14 unique hub genes demonstrated significant survival changes compared to the female modules (Fig. 6A, Supplementary Fig. 2A, Supplementary Table 3). These genes include ANK2, COL16A1, DAXX, DDX24, DLGAP5, ERCC5, FASN, HDAC6, IKBKB, LAMA2, NID1, PDGFRA, PPARG, SOX2. Among these, DLGAP5, SOX2, ERCC5, FASN, PPARG, DAXX, were upregulated, while HDAC6, IKBKB, DDX24, NID1, COL16A1, ANK2, LAMA2, PDGFRA were downregulated. Notably, the overexpression of PDGFRA, NID1, LAMA2, FASN, DLGAP5, DDX24, DAXX, ANK2 was associated with significantly decreased survival in male patients. Conversely, low expression of IKBKB, HDAC6, ERCC5, ATR in male samples correlated with poorer survival outcomes. In the female modules, there are three unique hub genes, RAD51C, COL5A2, COL22A1 was identified (Fig. 6B, Supplementary Table 3). All these genes were upregulated in female tumor samples and the overexpression of these genes was linked to poorer survival. For the genes common to both male and female clusters, two showed significant survival changes in males, which are ATR and PXDNL, and two in females, which are COL4A4 and E2F8. ATR, PXDNL and E2F8 were upregulated and COL4A4 was downregulated in tumor samples for both sexes. These findings underscore the distinct and overlapping roles of the hub genes in male and female PPI networks and their potential impact on patient survival.

**Figure 6.**
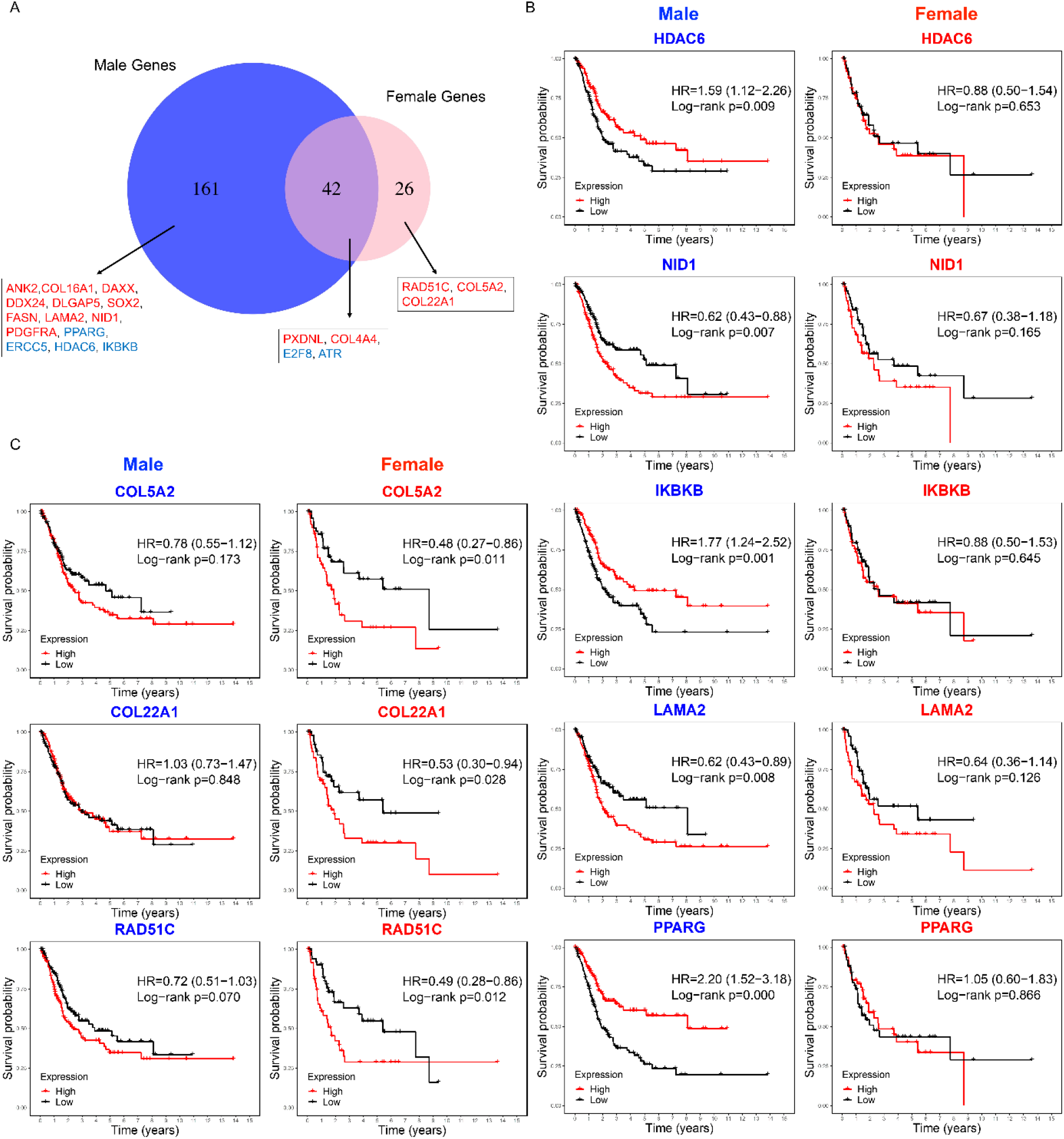
Sex-specific survival analysis for genes in the key modules of the two sexes in BLCA patients. (A) Venn diagram of genes in significant modules for both sexes. Genes identified as significant in survival analysis are indicated in the text boxes. Red and blue represent genes whose overexpression in all tumor samples is associated with poor or good survival outcomes, respectively. (B) top 5 genes significant for males only and (C) genes significant for females only (P < 0.05).

### 6. Immune cell filtration correlation analysis

We analyzed the abundance levels of the 22 immune cell types in female and male tumor samples using the CIBERSORT algorithm. Figure 7A illustrates the percentage distribution of immune cells in each sample, highlighting that macrophages M2, plasma cells, activated memory T cells, resting CD4+ memory T cells, CD8+ T cells, and follicular helper T cells constitute a large proportion (Supplementary Table 4). Violin plots revealed that activated mast cells, resting NK cells, plasma cells, and CD8+ T cells are slightly higher in male samples compared to the female samples (p < 0.05). To further investigate the difference between sex-specific and immune related hub genes and immune cells, we calculated the spearman correlation between each sex-specific hub gene and the remaining immune cell types for male and female tumor samples, respectively, with the five immune cell types with negligible proportions excluded. We compared the hub genes of male and female with the immune genes list from the ImmPort datasets to identify the immune related genes including DAXX, IKBKB, PDGFRA and PPARG, which are all male specific hub genes. As depicted in Figure 7B, significant correlations (p < 0.01) between gene expression and immune cell proportions to either male or female samples are marked with asterisk. DAXX demonstrated significant correlations with naïve B cells, M0 macrophages, resting mast cells, monocytes and activated CD4+ memory T cells in male while only has significant correlation with regulatory T cells in female tumor samples. This is the same case for IKBKB that showed more significant correlation with immune cells in males, including activated dendritic cells, M0 and M1 macrophages, monocytes, activated CD4+ memory T cells and regulatory T cells, in males but only one significant correlation with macrophages M0 in females. While PDGFRA showed unique significant correlation with resting CD4+ memory T cells and resting mast cells in females and male samples showed more significant correlation with the immune cells. The correlations for PPARG expression with the proportions of different immune cell are similar in males and females while males have unique significant correlation with activated CD4 memory T cells and monocytes. These findings suggested that these gender-specific hub genes may differentially regulate interactions with immune cells.

**Figure 7.**
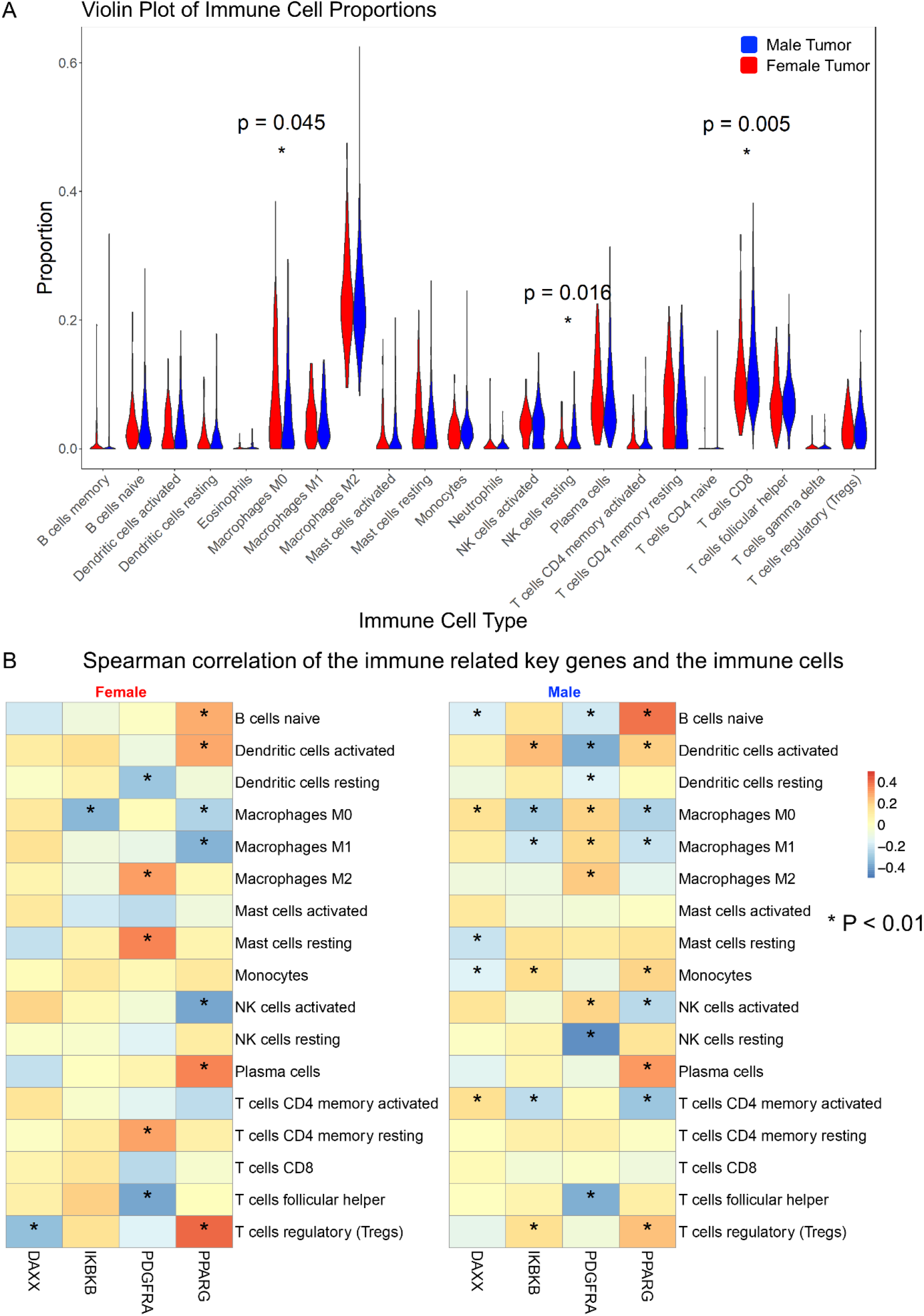
(A) Proportions of immune cells BLCA male and female samples. (B) Spearman correlation of the immune-related genes and the immune cells in female and male samples. (P<0.01)

## Discussions

Bladder cancer is a malignancy with high mortality rates and currently inadequate treatment options. A study in 2008 by Horstmann et al showed that the incidence of bladder cancer is higher in men than in women, with a male-to-female bladder cancer incidence ratio of 2.2:1, and men were diagnosed at a younger age compared to women, and although the histology of tumors did not differ between sexes, muscle-invasive tumors were more frequent in men (39.8% vs 34.5%) [17]. While sex differences in physiological bladder gene expression have been extensively studied in mice and humans, there is a paucity of knowledge regarding sex-dependent gene expression in human BLCA, which may have significant biological and medical implications.

In this study, we integrated mRNA profile datasets from TCGA and GTEx, along with whole-exome sequencing datasets from TCGA, using bioinformatic analysis methods to identify sex-associated differentially expressed genes with intermediate mutation frequency in BLCA tissues from male and female patients. In total, we identified 1,666 and 871 such genes in male and female patients, respectively. The PPI network analysis used these genes for males and females to identify significant key network modules. We identified four significant modules in males and two in females, respectively. There are 203 potential hub genes from the four male modules and 68 potential hub genes from the female modules. GO terms and KEGG pathway analysis for male and female hub genes showed highly similar enriched pathways, such as DNA repair, response to xenobiotic stimulus, p53 signaling pathway, PI3K-Akt signaling pathway, Fanconi anemia pathway, and ECM-receptor interaction, which are all critical cancer-related pathways [18][19][20][21][22][23]. This suggests that the two sexes likely employ highly similar regulation for tumorigenesis. Notably, we identified androgen receptor (AR) related pathways unique to male hub genes, while the Wnt signaling pathway was unique to female hub genes. AR signaling promotes bladder cancer development and progression, potentially explaining sex-specific differences in bladder cancer incidence and outcomes [24]. Shiota et al., showed that AR signaling contributes to tumor growth and drug resistance in bladder cancer cells, suggesting its potential as a therapeutic target [25]. AR signaling mediates CD8+ T cell exhaustion, contributing to sex differences in tumor aggressiveness, with male-biased expression driven by androgens impacting antitumor immunity [26]. Inhibition of the AR axis enhances T cell activity and improves the efficacy of immune checkpoint blockade therapies [27]. Furthermore, AR signaling promotes epithelial-mesenchymal transition (EMT) and metastasis in bladder cancer through the Wnt pathway and upregulation of the transcription factor Slug [28]. Androgen activation of Wnt/β-catenin signaling is linked to tumor progression, with AR promoting nuclear translocation and interaction with T-cell factor (TCF) in bladder cancer cells [29]. For the Wnt pathway unique for female hub genes, it is a key modulator of cellular proliferation and stem cell homeostasis. Pierzynski et al, has shown genetic variants in the Wnt signaling pathway have been identified as indicators of bladder cancer risk [30]. These observations suggest that the male specific hub genes uniquely enriched in AR signaling pathway and female specific hub genes uniquely enriched in Wnt signaling pathways may enhance sex differences in BLCA tumorigenesis and disease progression.

In survival analysis, we identified 14 male hub genes and three female hub genes related to the survival rate. Several genes function in the tumorigenesis of BLCA and have been shown to be potential prognostic biomarkers. DLGAP5 plays a critical role in cell cycle regulation, specifically in the mitotic phase, which is essential for cell proliferation. Its overexpression may lead to uncontrolled cell division, a hallmark of cancer. Rao et al,. has shown that high DLGAP5 expression is associated with poor prognosis [31]. The SOX2 gene encodes a transcription factor that plays a critical role in various biological processes, particularly in the maintenance of pluripotency and the regulation of development in stem cells. It has been shown that SOX2 is associated with stem-like properties, tumor aggressiveness, poor prognosis, and chemoresistance [32][33][34]. LAMA2 has been shown to significantly inhibit proliferation, weaken invasiveness, and promote apoptosis of BLCA cells [35]. High COL5A2 expression is associated with worse clinical outcomes, higher tumor grades, and poorer survival rates, making it a potential prognostic biomarker [36][37]. Interestingly, our study showed that DLGAP5, COL16A1, and LAMA2 are favorable prognostic biomarkers in males, while COL5A2 is favorable in females. We also identified other favorable prognostic biomarkers for males, such as FASN, PPARG, DAXX, ATR, HDAC6 and PDGFRA. High FASN expression in bladder cancer is related to immune cell infiltration and response to immune checkpoint inhibitors, and is associated with tumor progression and poor prognosis [38][39]. PPARG influences tumor growth, immune evasion, and cellular differentiation, making it a valuable target for therapeutic intervention [40][41][42][43]. PPARG activation through genetic alterations presents a valuable target for therapeutic intervention, potentially improving outcomes in bladder cancer treatment [44]. DAXX, a gene involved in chromatin remodeling, apoptosis regulation, and maintaining genomic stability has been shown that the overexpression of DAXX in various cancers has been linked to tumorigenesis, disease progression, and treatment resistance[45]. ATR plays a significant role in the DNA damage response and genomic stability in bladder cancer. Its expression correlates with tumor aggressiveness and resistance to therapy [45][46]. HDAC6 plays a critical role in the migration, invasion, and progression of bladder cancer. Targeting HDAC6 with specific inhibitors shows promise as a therapeutic strategy, particularly when combined with immunotherapies[47][48][49][50]. IKBKB plays a significant role in the progression of bladder cancer by regulating the NF-κB signaling pathway. It is a promising target for therapeutic intervention, particularly in combination with immunotherapy and other targeted treatments[51][52]. PDGFRA is expressed in various subtypes of bladder cancer and may play a role in tumor progression and metastasis. The presence of PDGFRA mutations and overexpression of its ligands suggest it could be a potential therapeutic target, especially in aggressive and metastatic bladder cancers [53][54]. ERCC5, NID1, and ANK2 are also male-favorable prognosis biomarkers, but there are no direct studies available for bladder cancer. This suggests further studies should be done for these genes related to male bladder cancer. For female-favorable prognosis biomarkers, COL5A2 was linked to essential pathways in cancer invasion, including cell adhesion and epithelial-mesenchymal transition. Several studies have demonstrated that the high expression of COL5A2 was associated with poorer prognosis[36][37][55]. No direct studies are available for the genes RAD51C and COL22A1, so further studies may be helpful to validate their potential role serving as prognosis biomarkers in female bladder cancer patients.

By comparing with known immune-related genes from the Immortal dataset, we identified four male hub genes overlapped with these immune-related genes, which are DAXX, IKBKB, PDGFRA, and PPARG. The spearman correlation analysis of the immune-related genes with various immune cells in male and female bladder cancer samples reveals noteworthy sex-specific differences. Males exhibited a higher number of significant correlations with different immune cells compared to females. The potential explanation for these observed differences lies in the influence of hormonal regulation, particularly by androgens. Androgens, such as testosterone, have been shown to modulate immune responses by influencing gene expression and signaling pathways. For instance, testosterone alters T-cell immunity by up-regulating Ptpn1, which inhibits IL-12 signaling in CD4 T cells, thereby affecting the activity of monocytes and NK cells [56]. DAXX (Death Domain-Associated Protein) has been studied to function as a negative androgen receptor coregulator through direct protein-protein interactions [57] playing a critical role in modulating immune responses. Its overexpression in various cancers has been linked to tumorigenesis and disease progression [45]. The suppression of DAXX expression sensitizes cells to stress-induced apoptosis [58]. Additionally, the significant positive correlations between DAXX and immune cells such as activated NK cells and monocytes in male bladder cancer samples suggest that DAXX might influence the immune landscape more prominently in males. IKBKB is a key regulator of the NF-κB signaling pathway, crucial for immune response regulation and inflammation. It has been demonstrated that androgen signaling can suppress T cell immunity via the NF-κB pathway, involving IKBKB, which might explain the more significant correlations observed in males [59]. This modulation can lead to enhanced immune cell interactions and influence cancer progression and response to therapy. PDGFRA is another gene with significant correlations to multiple immune cells, particularly more in males. PDGFRA’s role in regulating cell proliferation and survival can impact the tumor microenvironment and immune cell infiltration, contributing to the observed gender differences [60]. PPARG (Peroxisome proliferator-activated receptor gamma), a member of the nuclear receptor superfamily, has demonstrated bidirectional crosstalk with AR signaling pathways in human prostate cancer. PPARG ligands can either suppress or enhance AR signaling depending on the cancer’s response to castration, whereas AR activation reduces PPARG levels [61]. The expression and transcriptional activity of PPARG can be inhibited by AR in human prostate cancer cells [62]. This interaction with androgen signaling pathways can modulate immune cell functions and influence cancer progression. Although the interaction of PPARG with AR in bladder cancer remains uninvestigated, evidence suggested that PPARG affected bladder cancer development and progression by regulating metastasis, apoptosis, proliferation and reactive oxygen species (ROS) and lipid metabolism. These effects are likely mediated through PPARG-SIRT1 feedback loops, the PI3K-Akt signaling pathway, and the WNT/β-catenin signaling pathway [63]. The significant positive correlations between PPARG and monocytes and activated mast cells in both genders highlight its role in shaping the immune landscape in bladder cancer.

The present study has several limitations. First, clinical variables such as age, disease state, tumor purity, smoking status, and bladder cancer subtypes were not considered, potentially reducing the study’s observational power. Second, the limited number of normal tissue samples increases the sensitivity to potential confounding factors, which reduces detection power. Therefore, future studies will conduct rigorous analyses on larger patient cohorts with better control of clinical confounding variables. Third, our study focused solely on protein-coding genes and did not explore sex differences in the expression of non-coding genes, which may play critical roles in elucidating sex differences in bladder cancer. Finally, sex differences arising from non-RNA-based mechanisms, such as translational regulation, protein function, and post-translational modifications, need to be integrated to achieve a more comprehensive understanding of sex differences in bladder cancer in the future.

In conclusion, our comprehensive analysis elucidates sex-specific molecular mechanisms in bladder cancer, underscoring the critical need to consider sex differences in cancer research and therapeutic strategies. Our findings indicate that, while both sexes exhibit highly similar regulatory mechanisms for tumorigenesis, distinct pathways, such as androgen receptor (AR) signaling in males and Wnt signaling in females, play pivotal roles in the progression and prognosis of bladder cancer. We also identified 14 unique hub genes in males and 3 in females, with 4 of the male-specific immune-related hub genes demonstrating differential interactions with immune cells between sexes. These insights could pave the way for more personalized and effective therapeutic interventions tailored to male and female patients.

## Methods

### 1. Data collection

The clinical data and gene expression profiles of bladder cancer (BLCA) patients were obtained from the UCSC Xena database (https://xenabrowser.net/datapages/). In our analysis, we included a total of 303 tumor samples and 10 adjacent normal samples from male BLCA patients, and 108 tumor samples and 9 adjacent normal samples from female BLCA patients. Gene expression data from the Genotype-Tissue Expression (GTEx) project, comprising 14 male and 7 female samples, were downloaded and used in conjunction with the adjacent normal samples from TCGA as normal controls. Additionally, the Mutation Annotation Format (MAF) file containing somatic mutation information, analyzed from whole-exome sequencing data using VarScan2, was downloaded from TCGA database.

### 2. Screening of intermediate frequency mutation genes

The maftools package (version 2.14.0) was utilized to summarize the BLCA mutation data obtained from TCGA and to identify genes with intermediate mutation frequencies. The top 50 genes with highest mutation frequencies were displayed using a waterfall plot.

### 3. Differentially expressed gene identification

Protein-coding genes were selected, and low expressed genes (less than 20 counts in total across all samples) were removed. We yielded a total of 19,127 genes from the TCGA and GTEx samples. DEGs in tumor tissues compared to normal tissues for both male and female patients were identified using raw counts and analyzed with the DESeq2 package (version 1.38.3), applying an adjusted p-value threshold of < 0.05 for significance. The pheatmap R package (version 1.0.12) was utilized to generate the heatmap and volcano plot. DEGs that overlapped with intermediate frequency mutation genes, exhibiting a mutation frequency greater than 2%, were retained for further analysis.

### 4. GO and pathway enrichment analysis of DEGs

The Gene Ontology (G.O.) and the Kyoto Encyclopedia of Genes and Genomes (KEGG) pathway analyses were performed by clusterProfiler (Version 4.6.2), with adjusted p-value < 0.05 and gene number count ≥ 2. The top ten GO terms and KEGG pathways were visualized.

### 5. PPI network construction of DEGs and core genes identification

For the PPI analysis, STRING (Version:10.0, http://www.string-db.org/) was employed using all DEGs (adjusted p < 0.05; mf > 2%) in male and female samples, respectively, with a medium confidence 0.4 as the cutoff. Cytoscape (Version 3.2.0) was used to construct the network with the output from STRING. The most significant clustered modules within the PPI network were identified using the Cytoscape plug-in MCODE (Version 1.4.2, http://apps.cytoscape.org/apps/MCODE) method with a scoring threshold of ≥ 10. Validation was conducted using the CytoHubba plug-in (Version 0.1, https://apps.cytoscape.org/apps/cytohubba).

### 6. Key DEG expression validation and survival analysis of sex-biased genes

Gene expression data in transcripts per million (TPM) was downloaded from the UCSC Xena database. Tumor samples from male and female patients were retained for survival analysis. The R packages survival (version 3.6.4) and survminer (version 0.4.9) were utilized for the survival analysis, with gene expression data in TPM and overall survival information serving as inputs. Median mRNA expression levels were used to distinguish between high and low expression groups, and Kaplan-Meier curves were plotted. Statistical significance was assessed using the log-rank test.

### 7. Immune cell infiltration analysis

The deconvolution algorithm implemented in the R package CIBERSORT (version 0.1.0) was utilized to quantify the abundance levels of 22 immune cell subtypes based on the expression profiles of male and female samples, respectively. Samples with a significance threshold of P < 0.05 were included in further analyses. The Wilcoxon test was employed to compare the proportion differences in immune cell subtypes, and violin plots were generated to visualize the proportions in male and female tumor samples. After filtering out immune cell subtypes with proportions smaller than 0.5%, 17 immune cell subtypes remained. The Spearman correlation method was applied to assess the relationship between identified male immune-related hub genes and the 17 immune cell subtypes. Correlation heatmaps were generated using the R package pheatmap (version 1.0.12).

## Bibliography

[1] C. M. Lopes-Ramos, J. Quackenbush, and D. L. DeMeo, “Genome-Wide Sex and Gender Differences in Cancer,” Frontiers in Oncology, vol. 10, 2020, Accessed: Jul. 20, 2023. [Online]. Available: https://www.frontiersin.org/articles/10.3389/fonc.2020.597788

[2] J. B. Rubin et al., “Sex differences in cancer mechanisms,” Biology of Sex Differences, vol. 11, no. 1, p. 17, Apr. 2020, doi: 10.1186/s13293-020-00291-x.

[3] A. Dart, “Sexual dimorphism in cancer,” Nat Rev Cancer, vol. 20, no. 11, pp. 627–627, Nov. 2020, doi: 10.1038/s41568-020-00304-2.

[4] H. Li et al., “Connecting the mechanisms of tumor sex differences with cancer therapy,” Mol Cell Biochem, vol. 479, no. 2, pp. 213–231, Feb. 2024, doi: 10.1007/s11010-023-04723-1.

[5] S. S. Jackson et al., “Sex disparities in the incidence of 21 cancer types: Quantification of the contribution of risk factors,” Cancer, vol. 128, no. 19, pp. 3531–3540, 2022, doi: 10.1002/cncr.34390.

[6] B. Doshi, S. R. Athans, and A. Woloszynska, “Biological differences underlying sex and gender disparities in bladder cancer: current synopsis and future directions,” Oncogenesis, vol. 12, no. 1, Art. no. 1, Sep. 2023, doi: 10.1038/s41389-023-00489-9.

[7] C. M. Lam, Z. Li, D. Theodorescu, and X. Li, “Mechanism of Sex Differences in Bladder Cancer: Evident and Elusive Sex-biasing Factors,” Bladder Cancer, vol. 8, no. 3, pp. 241–254, doi: 10.3233/BLC-211658.

[8] C. H. Li, S. D. Prokopec, R. X. Sun, F. Yousif, N. Schmitz, and P. C. Boutros, “Sex differences in oncogenic mutational processes,” Nat Commun, vol. 11, no. 1, Art. no. 1, Aug. 2020, doi: 10.1038/s41467-020-17359-2.

[9] D. S. Chandrashekar et al., “UALCAN: A Portal for Facilitating Tumor Subgroup Gene Expression and Survival Analyses,” Neoplasia, vol. 19, no. 8, pp. 649–658, Aug. 2017, doi: 10.1016/j.neo.2017.05.002.

[10] J. Liu et al., “An Integrated TCGA Pan-Cancer Clinical Data Resource to Drive High-Quality Survival Outcome Analytics,” Cell, vol. 173, no. 2, pp. 400-416.e11, Apr. 2018, doi: 10.1016/j.cell.2018.02.052.

[11] L.-F. Zhao et al., “SDC: An integrated database for sex differences in cancer,” Computational and Structural Biotechnology Journal, vol. 20, pp. 1068–1076, Jan. 2022, doi: 10.1016/j.csbj.2022.02.023.

[12] J. J. de Jong et al., “Distribution of Molecular Subtypes in Muscle-invasive Bladder Cancer Is Driven by Sex-specific Differences,” European Urology Oncology, vol. 3, no. 4, pp. 420–423, Aug. 2020, doi: 10.1016/j.euo.2020.02.010.

[13] Q. Wang et al., “Unifying cancer and normal RNA sequencing data from different sources,” Sci Data, vol. 5, no. 1, p. 180061, Apr. 2018, doi: 10.1038/sdata.2018.61.

[14] W. Z. D. Zeng, B. S. Glicksberg, Y. Li, and B. Chen, “Selecting precise reference normal tissue samples for cancer research using a deep learning approach,” BMC Medical Genomics, vol. 12, no. 1, p. 21, Jan. 2019, doi: 10.1186/s12920-018-0463-6.

[15] M. S. Lawrence et al., “Discovery and saturation analysis of cancer genes across 21 tumour types,” Nature, vol. 505, no. 7484, pp. 495–501, Jan. 2014, doi: 10.1038/nature12912.

[16] A. G. Robertson et al., “Comprehensive Molecular Characterization of Muscle-Invasive Bladder Cancer,” Cell, vol. 171, no. 3, pp. 540-556.e25, Oct. 2017, doi: 10.1016/j.cell.2017.09.007.

[17] M. Horstmann, R. Witthuhn, M. Falk, and A. Stenzl, “Gender-specific differences in bladder cancer: a retrospective analysis,” Gend Med, vol. 5, no. 4, pp. 385–394, Dec. 2008, doi: 10.1016/j.genm.2008.11.002.

[18] T. Helleday, E. Petermann, C. Lundin, B. Hodgson, and R. A. Sharma, “DNA repair pathways as targets for cancer therapy,” Nat Rev Cancer, vol. 8, no. 3, pp. 193–204, Mar. 2008, doi: 10.1038/nrc2342.

[19] L. Wen and Z. Han, “Identification and validation of xenobiotic metabolism-associated prognostic signature based on five genes to evaluate immune microenvironment in colon cancer,” J Gastrointest Oncol, vol. 12, no. 6, pp. 2788–2802, Dec. 2021, doi: 10.21037/jgo-21-655.

[20] M. F. Ladelfa, M. F. Toledo, J. E. Laiseca, and M. Monte, “Interaction of p53 with tumor suppressive and oncogenic signaling pathways to control cellular reactive oxygen species production,” Antioxid Redox Signal, vol. 15, no. 6, pp. 1749–1761, Sep. 2011, doi: 10.1089/ars.2010.3652.

[21] A. Glaviano et al., “PI3K/AKT/mTOR signaling transduction pathway and targeted therapies in cancer,” Mol Cancer, vol. 22, no. 1, p. 138, Aug. 2023, doi: 10.1186/s12943-023-01827-6.

[22] W. Liu, A. Palovcak, F. Li, A. Zafar, F. Yuan, and Y. Zhang, “Fanconi anemia pathway as a prospective target for cancer intervention,” Cell Biosci, vol. 10, p. 39, 2020, doi: 10.1186/s13578-020-00401-7.

[23] M. Najafi, B. Farhood, and K. Mortezaee, “Extracellular matrix (ECM) stiffness and degradation as cancer drivers,” J Cell Biochem, vol. 120, no. 3, pp. 2782–2790, Mar. 2019, doi: 10.1002/jcb.27681.

[24] P. Li, J. Chen, and H. Miyamoto, “Androgen Receptor Signaling in Bladder Cancer,” Cancers (Basel), vol. 9, no. 2, p. 20, Feb. 2017, doi: 10.3390/cancers9020020.

[25] M. Shiota et al., “Androgen receptor signaling regulates cell growth and vulnerability to doxorubicin in bladder cancer,” J Urol, vol. 188, no. 1, pp. 276–286, Jul. 2012, doi: 10.1016/j.juro.2012.02.2554.

[26] H. Kwon et al., “Androgen conspires with the CD8+ T cell exhaustion program and contributes to sex bias in cancer,” Sci Immunol, vol. 7, no. 73, p. eabq2630, Jul. 2022, doi: 10.1126/sciimmunol.abq2630.

[27] T. Xiao et al., “Novel mechanisms of androgen receptor-centered transcriptional regulatory network in regulating CD8 +T cell exhaustion and sex bias in cancer,” The Journal of Immunology, vol. 210, no. 1_Supplement, p. 171.13-171.13, May 2023, doi: 10.4049/jimmunol.210.Supp.171.13.

[28] Y. Jing et al., “Activated androgen receptor promotes bladder cancer metastasis via Slug mediated epithelial-mesenchymal transition,” Cancer Letters, vol. 348, no. 1–2, pp. 135–145, Jun. 2014, doi: 10.1016/j.canlet.2014.03.018.

[29] Y. Li et al., “Androgen activates β-catenin signaling in bladder cancer cells,” Endocrine-Related Cancer, vol. 20, no. 3, pp. 293–304, Jun. 2013, doi: 10.1530/ERC-12-0328.

[30] J. A. Pierzynski et al., “Genetic Variants in the Wnt/β-Catenin Signaling Pathway as Indicators of Bladder Cancer Risk,” J Urol, vol. 194, no. 6, pp. 1771–1776, Dec. 2015, doi: 10.1016/j.juro.2015.07.032.

[31] X. Rao, H. Cao, Q. Yu, X. Ou, R. Deng, and J. Huang, “NEAT1/MALAT1/XIST/PKD--Hsa-Mir-101-3p--DLGAP5 Axis as a Novel Diagnostic and Prognostic Biomarker Associated With Immune Cell Infiltration in Bladder Cancer,” Front. Genet., vol. 13, p. 892535, Jul. 2022, doi: 10.3389/fgene.2022.892535.

[32] F. Zhu et al., “SOX2 Is a Marker for Stem-like Tumor Cells in Bladder Cancer,” Stem Cell Reports, vol. 9, no. 2, pp. 429–437, Aug. 2017, doi: 10.1016/j.stemcr.2017.07.004.

[33] J. Ruan et al., “Predictive value of Sox2 expression in transurethral resection specimens in patients with T1 bladder cancer,” Med Oncol, vol. 30, no. 1, p. 445, Mar. 2013, doi: 10.1007/s12032-012-0445-z.

[34] G. Chen, Y. Chen, R. Xu, G. Zhang, X. Zou, and G. Wu, “Impact of SOX2 function and regulation on therapy resistance in bladder cancer,” Front. Oncol., vol. 12, p. 1020675, Nov. 2022, doi: 10.3389/fonc.2022.1020675.

[35] Y. Jin, S. Huang, and Z. Wang, “Identify and validate RUNX2 and LAMA2 as novel prognostic signatures and correlate with immune infiltrates in bladder cancer,” Front. Oncol., vol. 13, p. 1191398, Jul. 2023, doi: 10.3389/fonc.2023.1191398.

[36] X.-Y. Meng et al., “The Role of COL5A2 in Patients With Muscle-Invasive Bladder Cancer: A Bioinformatics Analysis of Public Datasets Involving 787 Subjects and 29 Cell Lines,” Front. Oncol., vol. 8, p. 659, Jan. 2019, doi: 10.3389/fonc.2018.00659.

[37] X.-T. Zeng, X.-P. Liu, T.-Z. Liu, and X.-H. Wang, “The clinical significance of COL5A2 in patients with bladder cancer: A retrospective analysis of bladder cancer gene expression data,” Medicine, vol. 97, no. 10, p. e0091, Mar. 2018, doi: 10.1097/MD.0000000000010091.

[38] A. E. Abdelrahman, H. E. Rashed, E. Elkady, E. A. Elsebai, A. El-Azony, and I. Matar, “Fatty acid synthase, Her2/neu, and E2F1 as prognostic markers of progression in non-muscle invasive bladder cancer,” Annals of Diagnostic Pathology, vol. 39, pp. 42–52, Apr. 2019, doi: 10.1016/j.anndiagpath.2019.01.002.

[39] Q. Xiong et al., “Fatty Acid Synthase Is the Key Regulator of Fatty Acid Metabolism and Is Related to Immunotherapy in Bladder Cancer,” Front. Immunol., vol. 13, p. 836939, Mar. 2022, doi: 10.3389/fimmu.2022.836939.

[40] J. T. Goldstein et al., “Genomic Activation of PPARG Reveals a Candidate Therapeutic Axis in Bladder Cancer,” Cancer Research, vol. 77, no. 24, pp. 6987–6998, Dec. 2017, doi: 10.1158/0008-5472.CAN-17-1701.

[41] N. Rochel et al., “Recurrent activating mutations of PPARγ associated with luminal bladder tumors,” Nat Commun, vol. 10, no. 1, p. 253, Jan. 2019, doi: 10.1038/s41467-018-08157-y.

[42] C. Liu et al., “Pparg promotes differentiation and regulates mitochondrial gene expression in bladder epithelial cells,” Nat Commun, vol. 10, no. 1, p. 4589, Oct. 2019, doi: 10.1038/s41467-019-12332-0.

[43] T. Tate et al., “Pparg signaling controls bladder cancer subtype and immune exclusion,” Nat Commun, vol. 12, no. 1, p. 6160, Oct. 2021, doi: 10.1038/s41467-021-26421-6.

[44] D. J. Sanchez, R. Missiaen, N. Skuli, D. J. Steger, and M. C. Simon, “Cell-Intrinsic Tumorigenic Functions of PPARγ in Bladder Urothelial Carcinoma,” Mol Cancer Res, vol. 19, no. 4, pp. 598–611, Apr. 2021, doi: 10.1158/1541-7786.MCR-20-0189.

[45] I. Mahmud and D. Liao, “DAXX in cancer: phenomena, processes, mechanisms and regulation,” Nucleic Acids Research, vol. 47, no. 15, pp. 7734–7752, Sep. 2019, doi: 10.1093/nar/gkz634.

[46] T. Schepeler et al., “A high resolution genomic portrait of bladder cancer: correlation between genomic aberrations and the DNA damage response,” Oncogene, vol. 32, no. 31, pp. 3577–3586, Aug. 2013, doi: 10.1038/onc.2012.381.

[47] W. Chen, “HDAC6 and SIRT2 promote bladder cancer cell migration and invasion by targeting cortactin,” Oncol Rep, Nov. 2011, doi: 10.3892/or.2011.1553.

[48] X. S. Xu, L. Wang, J. Abrams, and G. Wang, “Histone deacetylases (HDACs) in XPC gene silencing and bladder cancer,” J Hematol Oncol, vol. 4, no. 1, p. 17, Dec. 2011, doi: 10.1186/1756-8722-4-17.

[49] B. Burke et al., “Inhibition of Histone Deacetylase (HDAC) Enhances Checkpoint Blockade Efficacy by Rendering Bladder Cancer Cells Visible for T Cell-Mediated Destruction,” Front. Oncol., vol. 10, p. 699, May 2020, doi: 10.3389/fonc.2020.00699.

[50] L. Rosik, G. Niegisch, U. Fischer, M. Jung, W. A. Schulz, and M. J. Hoffmann, “Limited efficacy of specific HDAC6 inhibition in urothelial cancer cells,” Cancer Biology & Therapy, vol. 15, no. 6, pp. 742– 757, Jun. 2014, doi: 10.4161/cbt.28469.

[51] J. Yang, D. Yuan, J. Li, S. Zheng, and B. Wang, “miR-186 downregulates protein phosphatase PPM1B in bladder cancer and mediates G1-S phase transition,” Tumor Biol., vol. 37, no. 4, pp. 4331– 4341, Apr. 2016, doi: 10.1007/s13277-015-4117-4.

[52] J. Wei, X. Zeng, L. Han, and Y. Huang, “The regulatory effects of polyporus polysaccharide on the nuclear factor kappa B signal pathway of bladder cancer cells stimulated by Bacillus Calmette-Guerin,” Chin. J. Integr. Med., vol. 17, no. 7, pp. 531–536, Jul. 2011, doi: 10.1007/s11655-010-0787-y.

[53] T. Terada, “Autopsy case of primary small cell carcinoma of the urinary bladder: KIT and PDGFRA expression and mutations,” Pathology International, vol. 59, no. 4, pp. 247–250, Apr. 2009, doi: 10.1111/j.1440-1827.2009.02358.x.

[54] N. Eliyakin, H. Postaci, Y. Baskin, and Z. Kozacioğlu, “Small Cell Carcinoma of the Urinary Bladder: KIT and PDGFRA Gene Mutations,” Rare Tumors, vol. 7, no. 4, pp. 154–156, Dec. 2015, doi: 10.4081/rt.2015.5982.

[55] M. Ingenwerth, P. Nyirády, B. Hadaschik, T. Szarvas, and H. Reis, “The Prognostic Value of Cytokeratin and Extracellular CollagenExpression in Urinary Bladder Cancer,” CMM, vol. 22, no. 10, pp. 941–949, Dec. 2022, doi: 10.2174/1566524021666210225100041.

[56] H. T. Kissick et al., “Androgens alter T-cell immunity by inhibiting T-helper 1 differentiation,” Proc. Natl. Acad. Sci. U.S.A., vol. 111, no. 27, pp. 9887–9892, Jul. 2014, doi: 10.1073/pnas.1402468111.

[57] D.-Y. Lin et al., “Negative Modulation of Androgen Receptor Transcriptional Activity by Daxx,” Molecular and Cellular Biology, vol. 24, no. 24, pp. 10529–10541, Dec. 2004, doi: 10.1128/MCB.24.24.10529-10541.2004.

[58] L.-Y. Chen and J. D. Chen, “Daxx Silencing Sensitizes Cells to Multiple Apoptotic Pathways,” Molecular and Cellular Biology, vol. 23, no. 20, pp. 7108–7121, Oct. 2003, doi: 10.1128/MCB.23.20.7108-7121.2003.

[59] X. Zhang et al., “Androgen Signaling Contributes to Sex Differences in Cancer by Inhibiting NF-κB Activation in T Cells and Suppressing Antitumor Immunity,” Cancer Research, vol. 83, no. 6, pp. 906– 921, Mar. 2023, doi: 10.1158/0008-5472.CAN-22-2405.

[60] M. Bartoschek and K. Pietras, “PDGF family function and prognostic value in tumor biology,” Biochemical and Biophysical Research Communications, vol. 503, no. 2, pp. 984–990, Sep. 2018, doi: 10.1016/j.bbrc.2018.06.106.

[61] E. Olokpa, P. E. Moss, and L. V. Stewart, “Crosstalk between the Androgen Receptor and PPAR Gamma Signaling Pathways in the Prostate,” PPAR Res, vol. 2017, p. 9456020, 2017, doi: 10.1155/2017/9456020.

[62] E. Olokpa, A. Bolden, and L. V. Stewart, “The Androgen Receptor Regulates PPARγ Expression and Activity in Human Prostate Cancer Cells,” J Cell Physiol, vol. 231, no. 12, pp. 2664–2672, Dec. 2016, doi: 10.1002/jcp.25368.

[63] S. Lv, W. Wang, H. Wang, Y. Zhu, and C. Lei, “PPARγ activation serves as therapeutic strategy against bladder cancer via inhibiting PI3K-Akt signaling pathway,” BMC Cancer, vol. 19, no. 1, p. 204, Mar. 2019, doi: 10.1186/s12885-019-5426-6.

