## Supplementary figures and images for "An Integrative Study to Investigate Sex-Specific Biomarkers in Bladder Cancer Patients"

### supplementary_figure1_300dpi.tif

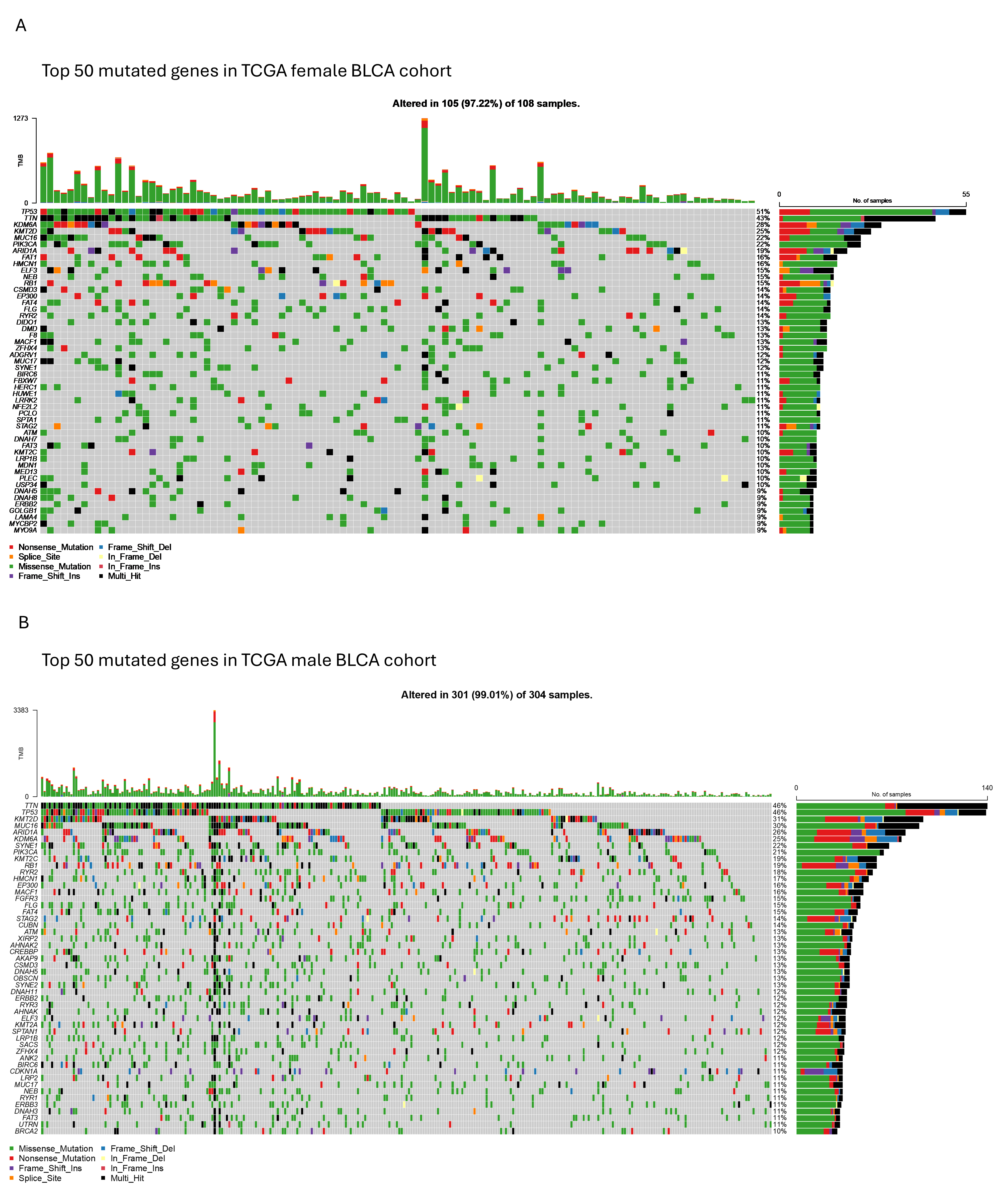

### supplementary_figure2_300dpi.tif

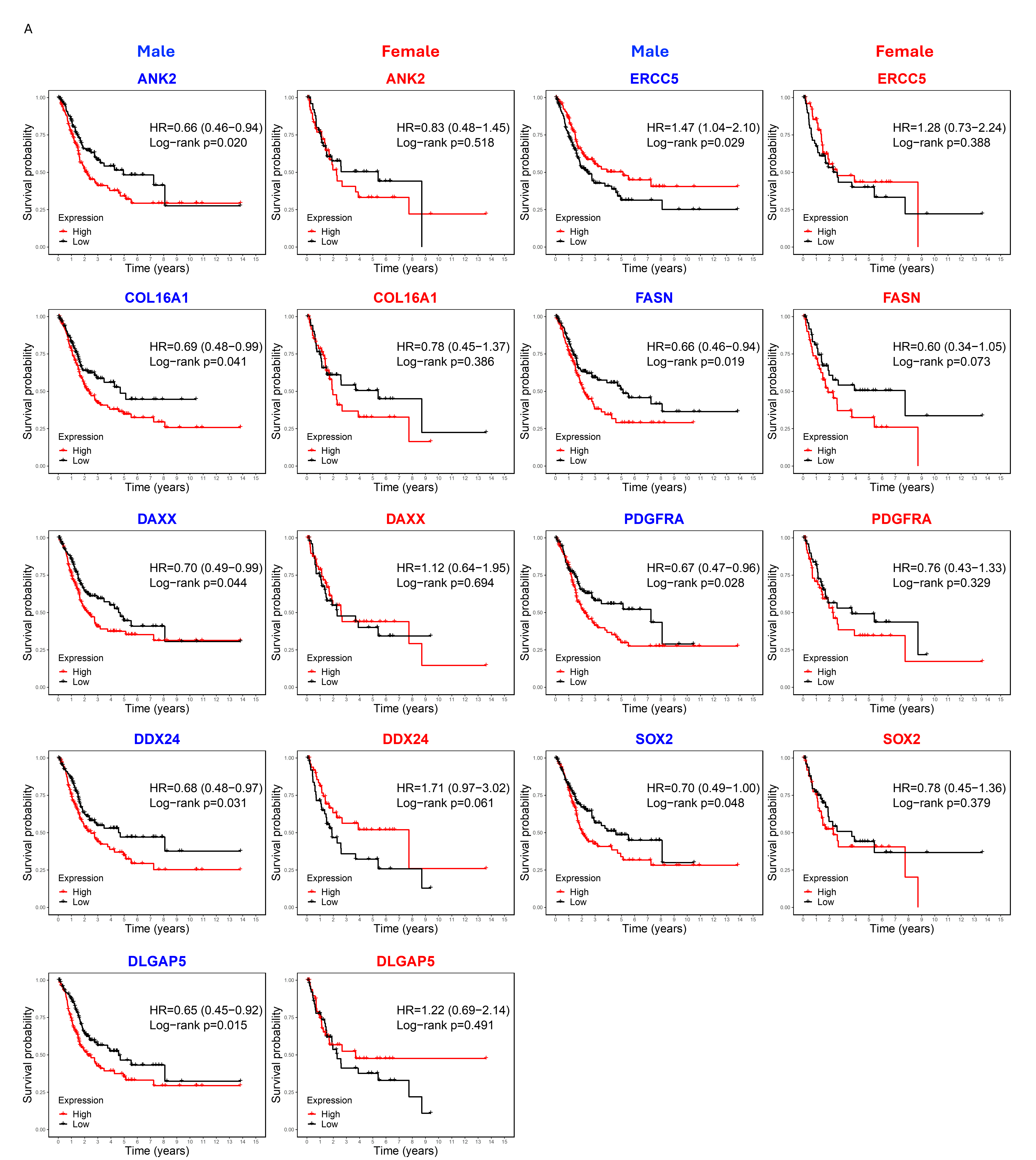
